# Percolation in the resting zebrafish habenula

**DOI:** 10.1101/481358

**Authors:** Suryadi, Ruey-Kuang Cheng, Suresh Jesuthasan, Lock Yue Chew

## Abstract

The habenula is an evolutionarily conserved structure of the vertebrate brain that is essential for behavioural flexibility and mood control. It is spontaneously active and is able to access diverse states when the animal is exposed to sensory stimuli or reward. Here we analyze two-photon calcium imaging time-series of the habenula of larval zebrafish and find that percolation occurs, indicating the presence of long-range spatial correlations within each side of the habenula, with percolation occurring independently in each side. On the other hand, the analysis of neuronal avalanches suggests that the system is subcritical, implying that the flexibility in its dynamics may result from other dynamical processes.

## Introduction

A defining feature of neural circuits is their flexibility. Although anatomical connectivity does not change rapidly, functional connectivity can change within seconds, enabling animals to quickly alter their behavior to deal with changing circumstances. This flexibility is made possible in part by the action of neuromodulators [1, 2, 3]. To ensure that functional connectivity is appropriate for a given condition, the release of neuromodulators is tied to a variety of factors, such as internal state of the animal, external cues, and perceived reward.

The habenula is an epithalamic structure that receives indirect input from several sensory systems [4, 5, 6], is stimulated by punishment or absence of an expected reward [7, 8, 9], and has activity that is dependent on the circadian clock [10] and stress levels [11]. It sends output to a number of structures, including the raphe [12] and the rostro-medial tegmental nucleus [13], which regulate the release of serotonin and dopamine respectively. Thus, the habenula is well positioned to coordinate release of broadly acting neuromodulators based on a variety of factors. The importance of the habenula is illustrated by a range of defects seen in animals with a lesioned habenula, including an inability to cope with changing circumstances [14] or to learn effectively [15, 16].

How does the habenula process information to efficiently influence behaviour? Calcium imaging in zebrafish indicates that neural activity in the habenula is sensitive to various features of sensory input. Different concentrations of odours [17], as well as wavelengths [18] or levels of ambient illumination [5] trigger different patterns of activity. This can be represented as trajectories in state space, with reproducible trajectories for each stimulus, indicating that sensory stimuli is represented by population activity. Additionally, habenula neurons constantly fire action potentials, even in the absence of sensory stimulation. This was demonstrated in rats using electrical recordings of brain slices, and has also been found in the zebrafish, using two-photon calcium imaging of tethered animals [19, 20], or CaMPARI-based labelling of freely swimming fish [21]. These features – constant activity and the ability to occupy multiple different states - raise the possibility that the habenula may be operating at criticality, which is known to confer benefits in information processing [22, 23, 24, 25, 26].

Here, using the zebrafish, which has an easily accessible epithalamus, we ask whether the habenula could be at a critical state in vivo. To evaluate this, we begin by investigating the statistical physics of both spatial clusters and subsequently neuronal avalanches, whose probability distributions should have the form of a power law in critical systems [22, 27, 28, 29]. In addition, we also examine the scaling laws and universal function, which are more rigorous hallmarks of criticality [27, 28, 29].

## 1 Materials and Methods

### 1.1 Ethical Statement

Experiments were carried out using protocols approved by the IACUC of Biopolis, Singapore (# 171215).

### 1.2 Microscopy and Image Processing

Calcium imaging was carried out on a Nikon A1R-MP two-photon system attached to a FN1 microscope with a 1.1 NA 25x water dipping objective. Fish expressing nuclear-localized GCaMP6f under the control of the *elavl3* promoter, i.e. *Tg(elavl3:H2B-GCaMP6f)jf7* [30], were immobilized in mivacurium, then embedded dorsal up in 2 percent agarose in a glass bottom dish (MatTek). The Ti-Sapphire laser was tuned to 930 nm, and images were collected at around 1 stack per second using a resonant scanner. A piezo-drive motor (Mad City Labs) was used to focus, with a step size of 5 *µ*m. A total of 35 recordings of around 10 minutes each was obtained in this manner.

The regions-of-interests (ROI) representing individual neurons and their fluorescence intensities were extracted using suite2p [31]. The corresponding Δ*F*/*F* values were then computed, which were then fed into the spike inference algorithm MLspike [32], using the parameters corresponding to the fluorescence indicator GCaMP6f.

### 1.3 Cluster Definition

Suite2p allows the extraction of not just the fluorescence, but also the pixel coordinates of each neuron. This allows us to construct a connectivity matrix for a graph based on our criteria for defining a neuronal connection, which is when neurons are adjacent (which we define to be true when two ROIs share at least one pixel). A cluster, then, is one consisting of connected neurons firing together at the same time instance.

### 1.4 Power Law Analysis

Power law distributions are distributions of the following form:

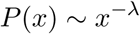

where *λ* represents the exponent of the power law distribution. This exponent is typically fit using maximum likelihood method. In this work, we bin the data logarithmically to obtain the density per bin prior to fitting the exponent, as it is well-known that logarithmic bins smooths the effect of noise [33].

We then adopt the method by [34] and compute the exponent *λ* by numerically solving a maximum likelihood equation:

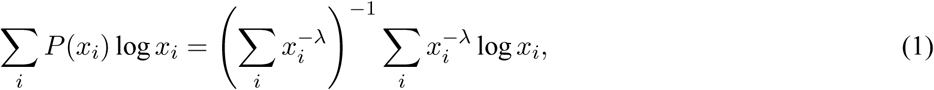

where *x*_*i*_ the *i*-th observation. This method is favored owing to its robustness over all values for the exponent. The KS statistic of each fit, which is a way to measure the distance between empirical data and its fit, was then computed via [35]:

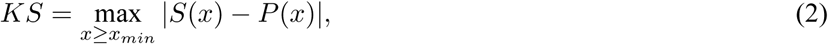

where *S*(*x*) and *P*(*x*) are the cumulative distribution function of the empirical data and fit distribution respectively, and *x*_*min*_ denotes the minimum *x* value above which lies the power law regime. The likelihood ratio test was also conducted in the manner proposed by the same work [35]: the parameters for candidate distributions, namely the exponential and lognormal distributions, were fit using standard maximum likelihood methods. The log-likelihood ratio for each pair of distributions (as well as their associated p-values) could then be computed, where the sign then determines which distribution is favored over the other.

As a sufficiently large sample size is required for the analysis of distributions, all power law analyses were conducted only when the number of data points are >100.

### 1.5 Shape collapse

Shape collapse was achieved by following an established workflow [29]. In theory, we expect that the average number of spikes *s*(*t, T*) for an avalanche of duration *T* at time *t* collapses to a universal parabolic shape when plotted as *s*(*t, T*)*T* ^−*γ*^ in the scale of normalized duration *t*/*T* for some exponent *γ*. We then optimized *γ* by minimizing the shape collapsed error, defined to be the mean variance divided by the range squared. To allow for such computations, *s*(*t, T*) for each duration *T* was linearly interpolated to 1000 points. Here we consider only avalanches lasting at least 5 time steps and occurring at least 50 times.

## 2 Results

### 2.1 Percolation

We performed volumetric calcium imaging of the habenula of larval zebrafish in the absence of stimulus (Fig 1). The recordings were then segmented into regions of interest (ROIs) corresponding to individual neurons, allowing us to extract their fluorescence intensities which we could then use for spike inference.

**Figure 1:**
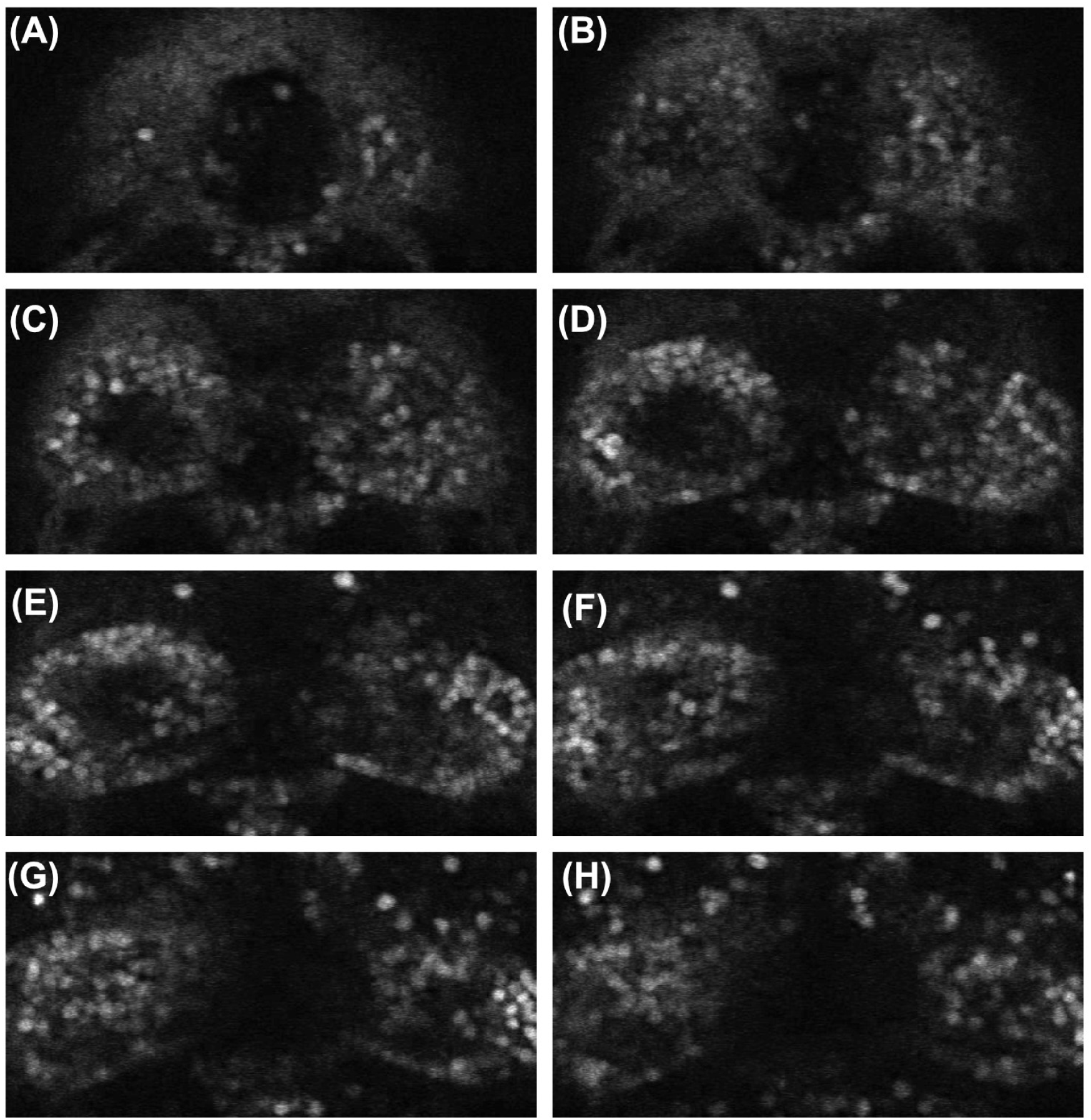
8 focal planes from one time point of a volumetric time-series recording of the habenula. Labels (A)-(I) denote planes 1-8 respectively.

We first analyzed the spatial statistics of clusters of neuronal spikes in the same manner as [36]. Here we consider an active neuron as belonging to a particular cluster if it is immediately adjacent to at least one neuron of that cluster. We then partitioned each frame of each recording, which has a certain fraction of active neurons *ρ*, into distinct clusters. Fig 2A shows a plot relating *ρ* and the number of clusters present. As percolation theory describes the transition from a regime of small, numerous clusters to one with few and large clusters, we expect the transition to occur near or at the point when the number of clusters no longer increases with increasing active units. As Fig 2A shows, we find this to occur at *ρ*_*c*_ = 0.15, noting that this is also the point with the largest spread and range in the number of clusters.

**Figure 2:**
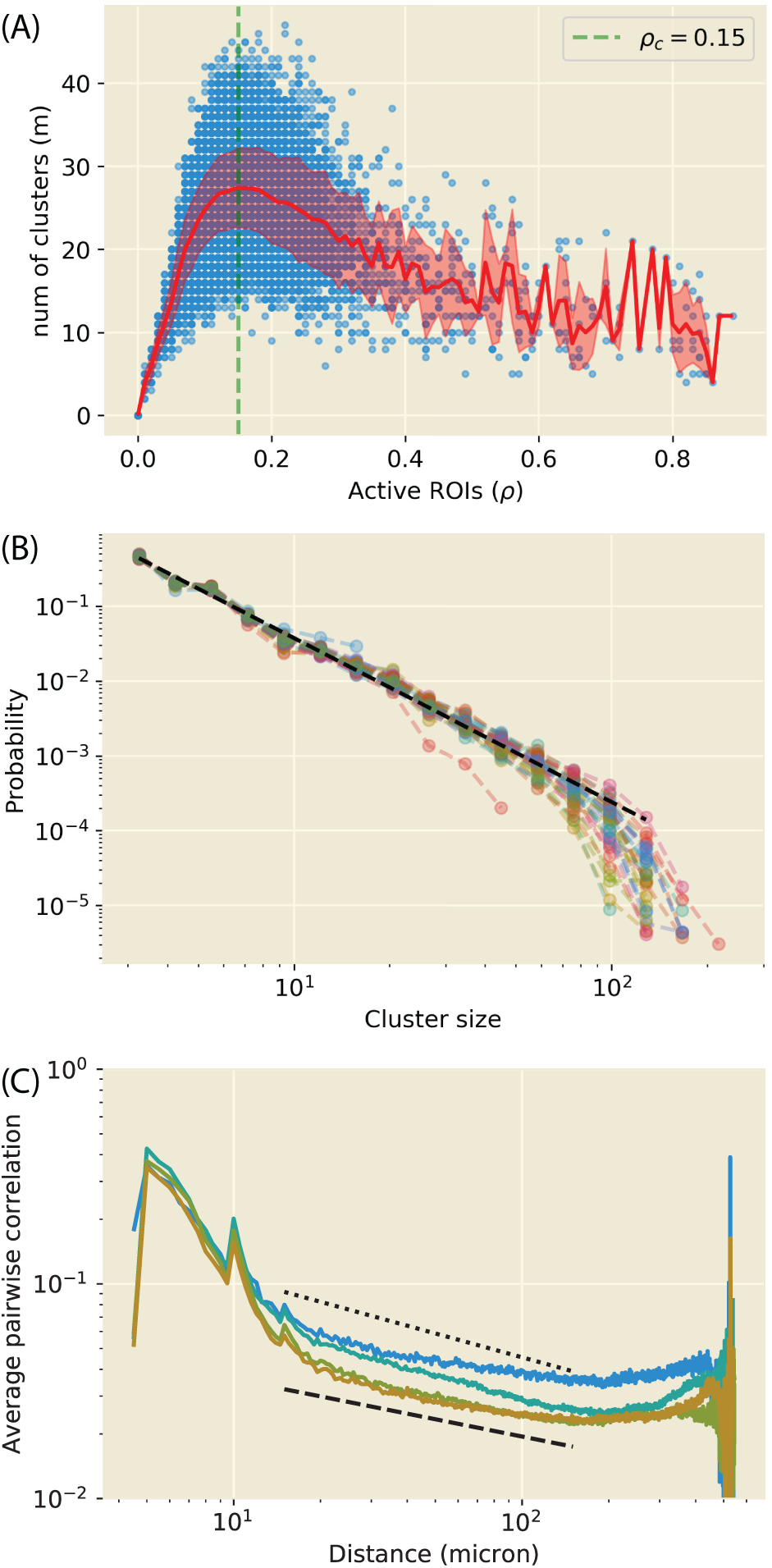
**(A)** The number of distinct clusters changes with the fraction of active ROIs (i.e. neurons) in the system. Each blue dot represents a data point (color intensity corresponding to the density of such points), with the red line and the surrounding pink region representing both the mean and standard error respectively. We see that the point of maximum number of clusters lies at *ρ*_*c*_ = 0.15, which is also the region with the greatest variability. **(B)** Cluster size distribution formed from clusters in the window *ρ* = *ρ*_*c*_ ± 0.02, which closely resembles a power law distribution. We obtain the exponent to be 2.20±0.09, which is consistent with the theoretical prediction at 2.19 (dashed line). **(C)** Pairwise correlation function for each pair of neurons in the dorsal habenula aggregated into four plots for each fish, with a power law-like regime as indicated. Three plots in the figure have their exponents computed to be *η* = 0.27 ± 0.03 (dashed plot), with another plot observed to have a significantly different exponent *η* = 0.37.

By collecting clusters near the obtained transition point, the cluster sizes, defined as the number of active neurons in a cluster, form a power law distribution (Fig 2B). Specifically, we considered only the clusters observed when *ρ* = *ρ*_*c*_ ± Δ where *ρ*_*c*_ = 0.15 and Δ = 0.02. We then fit the distribution using maximum likelihood methods to obtain the power law exponent, which we obtain to be 2.20 ± 0.09 (KS statistic = 0.027 ± 0.005). This value is consistent with the theoretical result of 2.19 for a 3D percolation process [37]. Likelihood ratio tests indicate that all cluster size distributions are likelier to be power law distributed than exponential or lognormal (*p* < 0.01). These results indicate that percolation dynamics occurs locally in the zebrafish habenula as well.

We then plotted the average pairwise correlation between neurons as a function of Euclidean distance (Fig 2C). As would be expected from a percolating system, we observed a power law decay in the correlation function as well, suggesting long-range spatial correlation between neurons. The exponents are obtained by least squares to be *η* = 0.27 ± 0.03 for three of the plots, with one significant deviation at *η* = 0.37. Regardless, the order of magnitude of these exponents is consistent with what has been found for the whole zebrafish brain [36]. Interestingly, however, we observed that the power law regime spans only up to the scale of any one side of the habenula (the diameter of each side was measured to be around 300 microns), beyond which the power law regime is no longer observable. This suggests that while each side of the habenula may be percolating, they may operate independently of each other.

### 2.2 Subcritical Avalanches

We then proceeded to investigate the spatiotemporal dynamics of the dorsal habenula to determine if the spatial correlation extends into the spatiotemporal domain as well. This was done by analyzing neuronal avalanches, which are cascades of neuronal firings propagating in both space and time. In particular, we considered these cascades to occur over connected neurons, and in the absence of finer connection information, only adjacent neurons are considered to be connected. Hence, if a neuron fires at the same time as a connected neuron, or if it fires after a connected neuron fired in a previous time step, these firings are taken to belong to the same avalanche. Any other avalanches are considered to be separate and hence analyzed separately and independently.

In this study, any active neuron connected to a member of an avalanche either in the current or previous time step was considered to also be a member of the same avalanche. An avalanche was considered to have been terminated if it reached a time step when no neuron within or connected to the avalanche fired. We then computed the avalanche size (the total number of firings over the course of an avalanche) and duration, as shown in Fig 3A-B.

**Figure 3:**
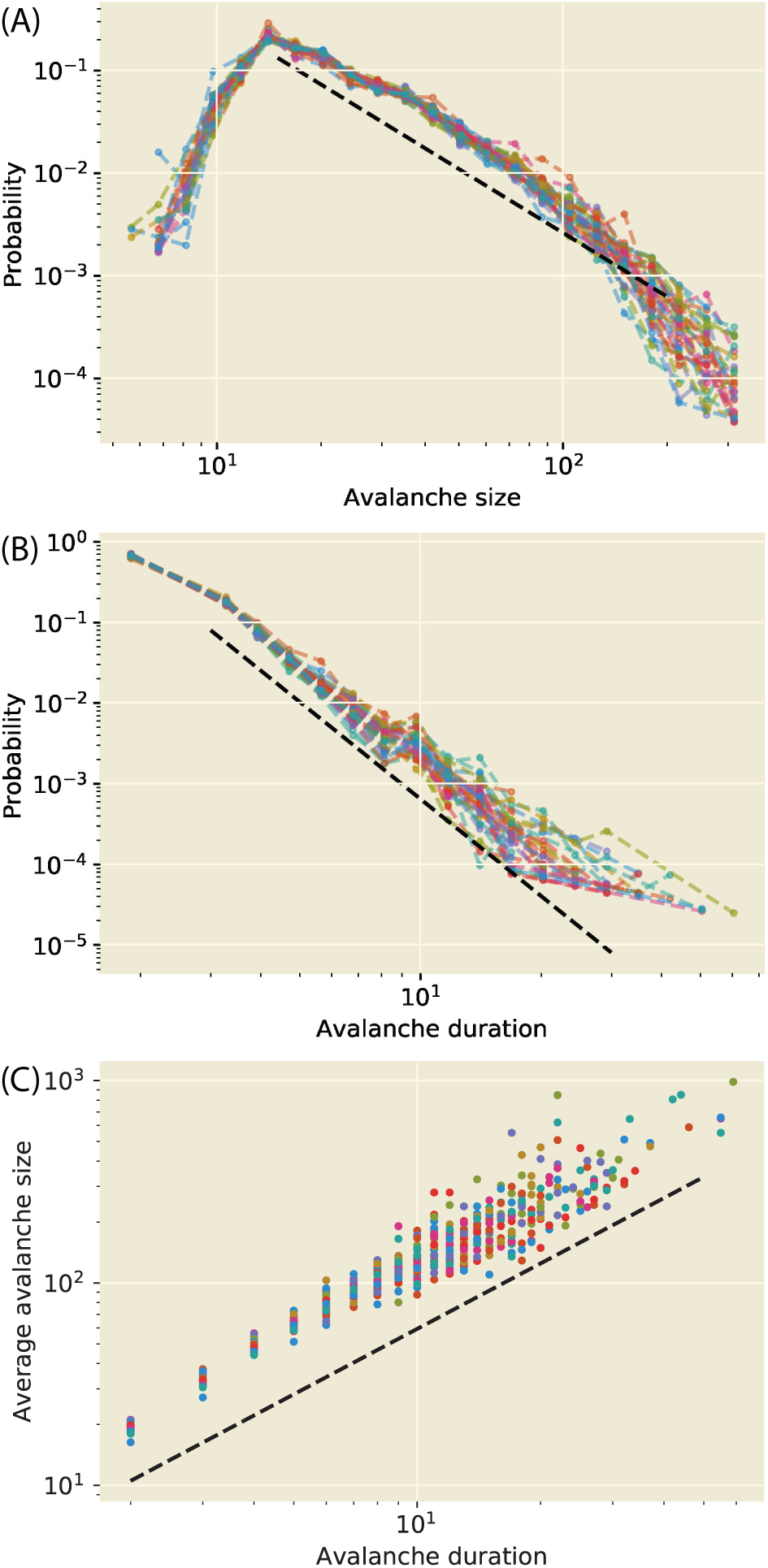
**(A)** Avalanche size distributions for each recording, with the dashed line representing the mean exponent, obtained to be 2.07 ± 0.22. We observe a sub-power law regime towards the right end of the plot, which suggests sub-criticality. **(B)** Avalanche duration distributions for each recording, with the dashed line representing the mean exponent of 4.00 ± 0.36. Unlike the avalanche size distribution, the duration distribution retains its power law regime. **(C)** The average avalanche size as a function of avalanche duration. Being a power function and not a distribution, the fitting is done not with maximum likelihood methods but with a standard least squares computed in log-log space, through which we obtain *β* = 1.07 ± 0.07 (dashed line).

For the case of a critical system, the avalanche size *s* and duration *T* distributions are power law distributed with well-defined exponents (denoted *τ* and *α* for avalanche size and duration respectively):

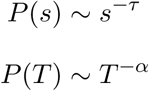

Fig 3A-B show the avalanche size and duration distributions respectively, which are binned logarithmically for both better visualization and fitting for the exponent [33]. We obtain *τ* = 2.07 ± 0.22 and *α* = 4.00 ± 0.36 (KS statistics = 0.029 ± 0.005 and 0.016 ± 0.006 respectively). Likelihood ratio tests indicate that both distributions are more likely to be power law than exponential or log-normal distributed (all distributions have *p* < 0.01 except for one size distribution).

A critical system exhibits self-similarity, which in this case also manifests in the temporal development of the avalanches, we expect avalanches of different durations to develop in a similar manner, differing only in their scales. In other words, with proper rescaling, the temporal development of the avalanches should collapse to the same shape (see Fig 4). The procedure for this collapse depends on a free parameter *γ* which we vary to minimize the average distance between the curves for each avalanche. This parameter *γ* is in fact closely related to a function which we can construct from the avalanche size *s* and duration *T* distributions, which is expected to form a power function with the exponent valued at *β* = 1 − *γ*:

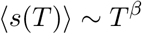

**Figure 4:**
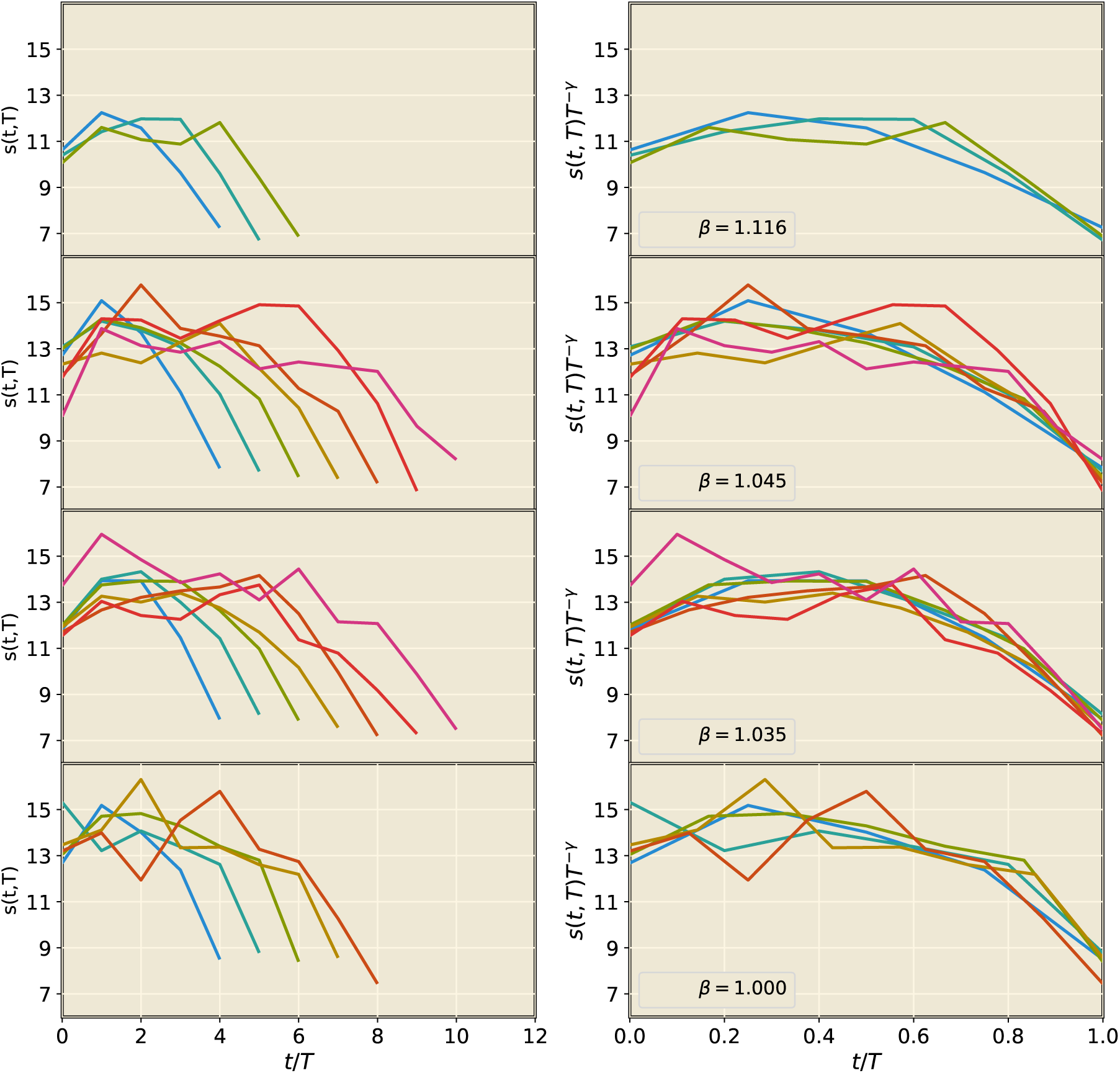
The average temporal profile of avalanches (left column) and the subsequent shape collapse (right column) for each fish in each row, which describes the average number of neurons *s*(*t, T*) firing at a particular time step (t) of an avalanche of a total duration *T*. Each curve averages over all avalanches of the same total duration. The critical exponent *β* can be transformed from the variational parameter for the shape collapse *γ* via *β* = 1 − *γ*, which are indicated in the plots. Averaging these four values yield *β* = 1.05 ± 0.04.

The plot of the function is given in Fig 3C, from which we obtain the exponent *β* to be 1.07 ± 0.07, which we see to closely agree with the *β* values obtained from shape collapse (Fig 4), which is on average 1.05 ± 0.04.

The three exponents *τ, α, β* are also known to satisfy a particular scaling relation in critical neuronal systems, namely:

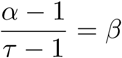

We see, however, that the scaling relation is not satisfied, despite the successful shape collapse and the agreement of the exponent *β*. Based on what we observe from the avalanche size distributions, which tapers off towards the right end leading to a sub-power law regime, it is likely that the zebrafish habenula is subcritical at rest. This is not in contradiction with other results as it is known that a non-critical system can nevertheless exhibit shape collapse [38].

## Discussion

In this article we have demonstrated the presence of percolation in the habenula of the zebrafish, consistent with what has been discovered for the entire zebrafish brain as a whole [36]. While specific to the habenula, we arrive at this conclusion also through a spike inference algorithm [32] instead of directly inferring spikes by thresholding the fluorescence intensity [36].

Upon analyzing the pairwise correlation function for the four datasets we observe two distinct behaviors for the power law regime. We saw that the four datasets can be divided into two groups with distinct power law exponents. We note that this may potentially reflect differences in mode of activity of the habenula, or simply be a variation due to the fish itself, which we are unable to determine at this point in time. Nevertheless, the scale invariant behavior of the function suggests long-range spatial correlations among the neurons in the habenula. More importantly, the scale invariant regime extends only up to the scale of any one side of the habenula, which suggests that percolation occurs independently in each side.

The presence of long-range spatial correlation suggests that the temporal aspect may display critical behavior as well. Despite so, our analysis suggests that the habenula are subcritical, i.e. the spatiotemporal neuronal dynamics are less active than the critical state.

In the absence of criticality, then, how could the habenula achieve its flexibility in terms of accessing different modes of activity? Two possibilities come to mind; the first of which is that the habenula operates not at, but near criticality, as has been suggested in literature [39]. Being near criticality may also explain why we observe the other telltale signs of criticality (including percolation), while failing only the more stringent properties (such as the scaling relation).

Another possibility lies in our fundamental assumption, which is that only adjacent cells are taken to be connected. While it is known that gap junction (which mediates communication among adjacent neurons) are present in habenula neurons [40], it could be the case that long-range connections play an important role in the development and transmission of avalanches, i.e. the habenula is actually critical, but we perceive it to be otherwise due to the omission of long-range connections. In such a situation, one large (and long-lasting) avalanche would be perceived as multiple smaller (and potentially shorter) avalanches, causing the probability mass in both distributions to bias towards the left side. Hence, it could explain the sub-power law regime of large avalanches in the size distribution, as well as the large magnitude of the exponent *α* in the duration distribution.

This work demonstrates that with our current available knowledge, while the zebrafish habenula displays percolation, its spatiotemporal behavior appears subcritical, which could mean either that its flexibility is a result of some other dynamical process, or that it is fundamentally critical when at rest, but that this underlying criticality is hidden from sight unless we include long-range connection information. A map of the habenula connectome could shed more light on the matter.

## Author Contributions

S and LC performed data analysis while RC and SJ performed two-photon imaging. The project was designed by SJ and LC.

## Acknowledgments

We would like to thank Dr. Mahathi Ramaswamy, Dr. He Chong, and Dr. Chua Khi Pin for the helpful discussions and prior explorations leading to this work.

